# Reconstructing the formation of Hmong-Mien genetic fine-structure

**DOI:** 10.1101/2022.11.23.517530

**Authors:** Zi-Yang Xia, Xingcai Chen, Chuan-Chao Wang, Qiongying Deng

## Abstract

The linguistic, historical, and subsistent uniqueness of Hmong-Mien (HM) speakers offers a wonderful opportunity to investigate how these factors impact the genetic structure. Nevertheless, the genetic differentiation among HM-speaking populations and the formation process behind are far from well characterized in previous studies. Here, we generated genome-wide data from 67 Yao ethnicity samples and analyzed them together with published data, particularly by leveraging haplotype-based methods. We identify that the fine-scale genetic substructure of HM-speaking populations corresponds better to linguistic classification than to geography, while the parallel of serial founder events and language differentiations can be found in West Hmongic speakers. Multiple lines of evidence indicate that ~500-year-old GaoHuaHua individuals are most closely related to West Hmongic-speaking Bunu. The excessive level of the genetic bottleneck of HM speakers, especially Bunu, is in agreement with their long-term practice of slash-and-burn agriculture. The inferred admixture dates in most of the HM-speaking populations overlap the reign of the Ming dynasty (1368 – 1644 CE). Besides the common genetic origin of HM speakers, their external ancestry majorly comes from neighboring Han Chinese and Kra-Dai speakers in South China. Conclusively, our analysis reveals the recent isolation and admixture events that contribute to the fine-scale genetic formation of present-day HM-speaking populations underrepresented in previous studies.

## Introduction

Consisting of ~40 different languages and dialects (Hammarström *et al*., 2022), the Hmong-Mien (HM) language family is currently spoken by ~6 million people living in the vast mountainous area of South China and Mainland Southeast Asia (Ratliff, 2010) [*Ratliff, p3*]. As suggested by historical linguistics, the age of the proto-HM language is ~500 BCE and is supposed to be originally spoken in the middle Yangtze Basin (Ratliff, 2021). According to historical records, the suggested precursors of the HM speakers, *Wŭxī Mán* (五溪蠻, literally “*Mán* of the Five Streams”), lived in the Wŭlíng Mountains (武陵山脈) between Hunan and Guizhou of Southwest China ~100-500 CE (Mao and Meng, 1986) [*p2*]. During the Tang (618-907 CE) and Song (960-1279 CE) dynasties, ethnonyms identical to present-day exonyms of HM-speaking groups, like Shē (畬/畲) and Yáo (徭/瑶), started to appear in Chinese historical documents (Zeng, 2005, Litzinger, 1995). Uprisings in Guizhou and Guangxi of Southwest China led by HM-speaking Miao and Yao people became more frequent during the reign of Ming (1368-1644 CE) and Qing (1644-1911 CE) dynasties (Scott, 2009) [*p137-140*], which was followed by the migration of some HM-speaking groups to Mainland Southeast Asia ~1800 CE (Kutanan *et al*., 2021).

Previous genomic studies have provided many intriguing insights into the population history of HM speakers, such as their shared genetic origin and admixture events during their past (Huang *et al*., 2022, Xia *et al*., 2019, Liu *et al*., 2020, Kutanan *et al*., 2021, Wang *et al*., 2021b, Yang *et al*., 2022). However, these studies focus more on the external genetic relationship of HM speakers to other groups (e.g., Sino-Tibetan and Kra-Dai speakers), while the formation history of genetic substructure within the HM speakers is obscure. For example, although the ~500-year-old GaoHuaHua population shares a similar genetic profile with the modern HM-speaking population (Wang *et al*., 2021b), it still remains unclear whether and with which modern HM-speaking populations they are more closely genetically related. Besides, haplotype-based methods, especially chromosome painting-based ones, have hardly ever been applied in previous studies, which have been shown to have larger power to identify fine-scale genetic history than allele frequency-based ones (Lawson *et al*., 2012, Leslie *et al*., 2015). The haplotype-based methods could enable us to investigate previously unresolved aspects of the genetic history of HM-speaking populations, such as exact sources of non-HM ancestries and the relationship between fine-scale genetic structure and language classifications.

In cultural anthropology, HM-speaking communities have often been collectively studied together with other ethnolinguistic groups in Southeast Asian mainland massif, also known as *Zomia* (Scott, 2009) [*preface, ix*]. Historically, the population in Zomia, including the majority of HM speakers, fell outside of the direct administration of the Sinicized and Indianized monarchies centralized at low elevations, where taxes and corvée labors were usually obligatory for the subjects (Scott, 2009) [*p13, p19, p116*]. In contrast to concentrated grain production (e.g., wet rice farming) practiced in the lowlands (Scott, 2009) [*p13*], HM-speaking Miao and Yao people historically practiced shifting and slash-and-burn agriculture in mountains regions [*p194-195*] (Scott, 2009). In fact, the exonyms of HM-speaking She (literally “slash-and-burn”) and Yao (originally *Mòyáo*/莫徭, “no corvée labor”) come from their historical practice of slash-and-burn agriculture and stateless, respectively (Zeng, 2005, Ozawa, 2000). Many recent genomic studies have focused on the populations historically marginalized from the state system in Europe (e.g., Roma) (Font-Porterias *et al*., 2019) and Africa (e.g., Ari Blacksmiths) (Van Dorp *et al*., 2015). In comparison, the genetic consequence of the stateless social practices of the population in Zomia, especially HM speakers, has never been investigated previously. Besides the historically documented Sinicization of HM speakers, anthropologists have also proposed an undocumented transition of Han Chinese into HM-speaking communities (Scott, 2009) [*p125-126*]. Such theories are also promising to be assessed by using genetic data.

It is indisputable that the HM language family is formed by two major branches: Hmongic and Mienic (Ratliff, 2021). In China, all the HM speakers are categorized into one of the following three officially recognized ethnicities: Miao, Yao, and She (Wang, 1983b, Mao *et al*., 1982, Mao and Meng, 1986). Yao is linguistically heterogeneous, including all the Mienic-speaking groups in China (except for Kin Mun in Hainan), Hmongic-speaking Bunu, Pahng, and Hmnai, Kra-Dai-speaking Lakkja, and currently Chinese-speaking Pingdi Yao (literally ‘Yao in plains’) (Simons and Fennig, 2017, Mao *et al*., 1982) [*p5-9*]. Guangxi has the largest population and richest linguistic diversity of the Yao ethnicity (Mao *et al*., 1982), but Yao populations in Guangxi have never been covered in previous genome-wide studies.

To overcome the aforementioned limitations in previous studies, we carried out this comprehensive investigation on the genetic differentiation within the HM speakers and the demographic and ancestral factors behind its formation, i.e., population isolation and admixture. We generated genome-wide SNP genotyping data from 67 Yao individuals from five Yao autonomous counties in Guangxi, whose geographic (Figure 1A) and linguistic (Figure 1B) information are shown in Figure 1. We addressed the three major issues throughout our study: What is the internal genetic substructure within the HM speakers, and how does it relate to ethnic identities (i.e., ethnic labels based on autonyms) and linguistic classifications? How does the social and cultural practice (e.g., shifting farming) in the HM history influence the pattern of population sizes across the HM-speaking populations? What is genetic interaction between HM and non-HM groups suggested by the inference of historical admixture events?

**Figure 1.**
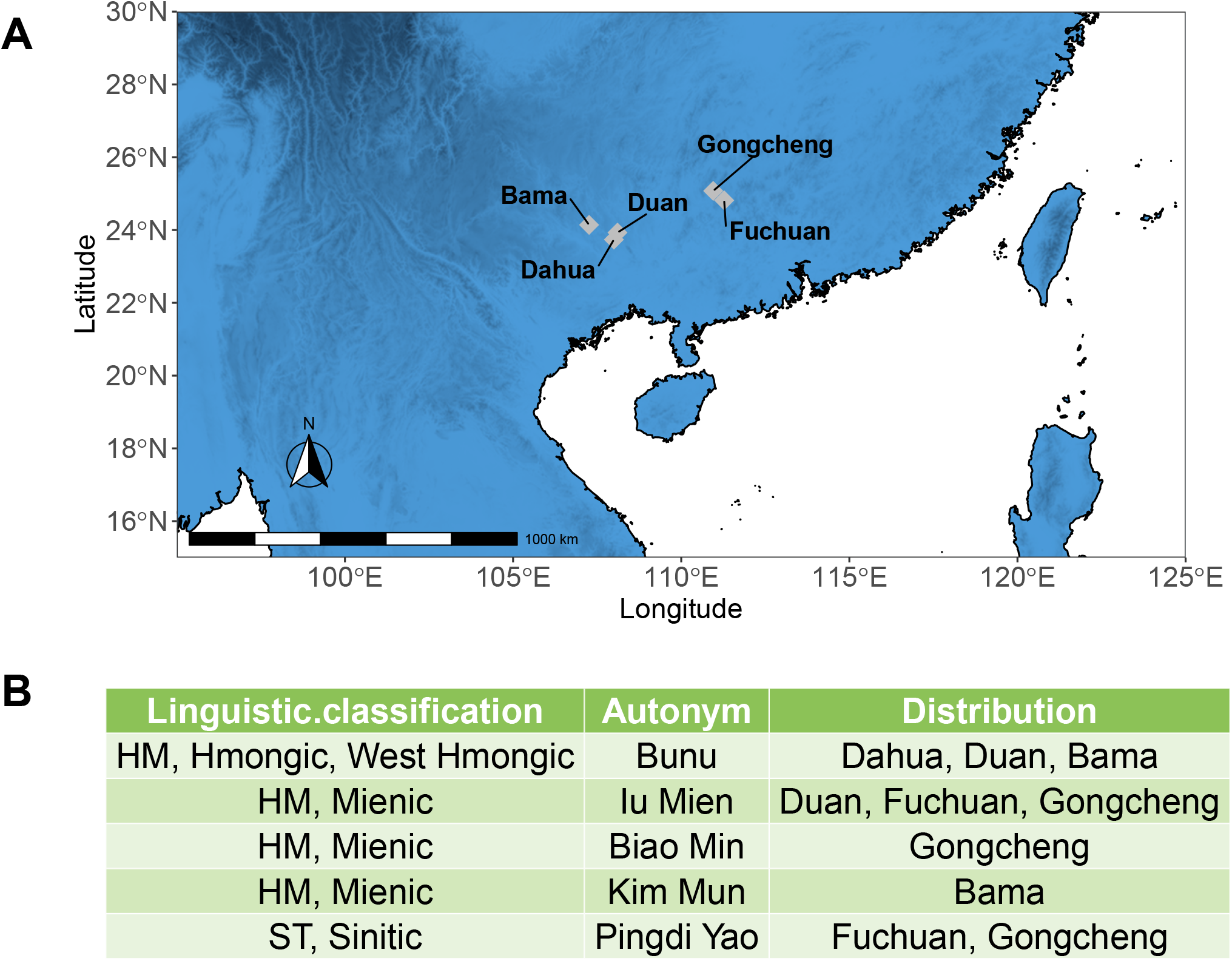
Geographic and linguistic information of newly reported samples. (A) Locations of all the five Yao autonomous counties where the individuals were sampled. (B) Languages spoken in all the five counties by people with Yao ethnicity and corresponding autonyms, according to Mao *et al*. (1982) and Meng (2001).

## Results

### An overview of population structure

We initially performed principal component analysis (PCA) to investigate the genetic variability of HM speakers in the contexts of southern East Asian populations (Figure 2), where ancient DNA (aDNA) samples were projected. In agreement with the previously reported language family-associated genetic structure of southern East Asians (Huang *et al*., 2022), HM speakers, including the newly reported individuals, form a distinct gradient apart from other southern East Asian populations (Figure 2). Likewise, ~500-year-old GaoHuaHua individuals are positioned together with present-day HM speakers, separating from other ancient genomes from South China. We also confirmed this genetic structure in ADMIXTURE, as HM speakers and GaoHuaHua share a distinct genetic component when K = 10 (SI Figure 1).

**Figure 2.**
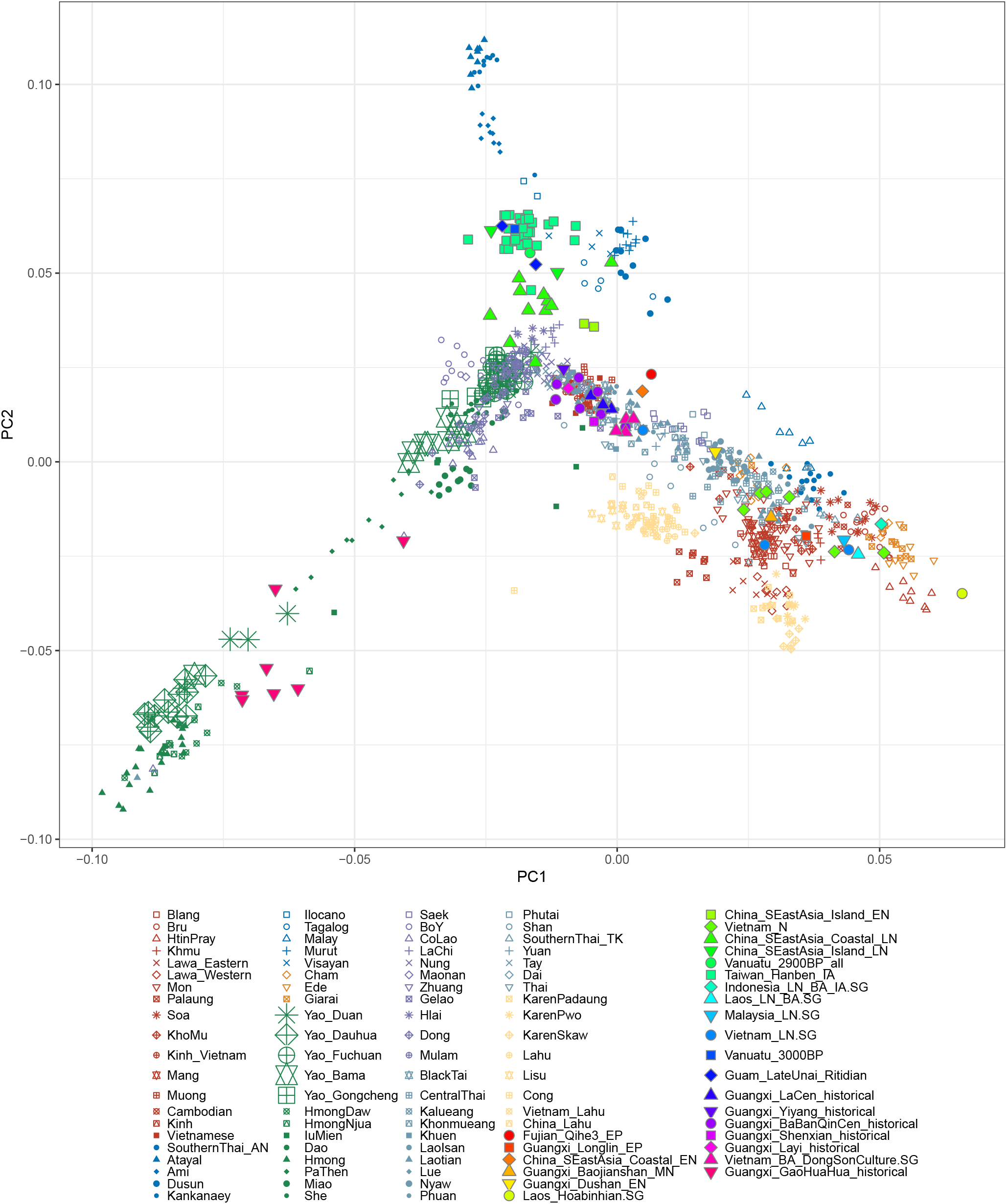
PCA of HM-speaking populations in the context of southern East Asians.

### Haplotype-based clustering

To further characterize the genetic substructure within the HM speakers, we applied haplotype-based ChromoPainter and fineSTRUCTURE (Figure 3) exclusively for HM-speaking individuals (Lawson *et al*., 2012). ChromoPainter infers each phased genotype as a mosaic of other most closely related genotypes, while fineSTRUCTURE leverages this haplotype-sharing pattern to cluster individual genotypes into a dendrogram. In general, individuals with the same autonym and linguistic affiliation tend to cluster together, suggesting that the marital practice might be more frequent in individuals with shared ethnic identities. Particularly, Yao from Du’an and Bama respectively group into two distinct genetic clusters (‘Bunu’ and ‘Mienic3’ for Yao_Duan, ‘Mienic1’ and ‘Bunu’ for Yao_Bama, respectively, Figure 3), consistent with the fact that both Hmongic (Bunu) and Mienic (Iu Mien and Kim Mun, respectively) languages are distributed in both localities (Figure 1B).

**Figure 3.**
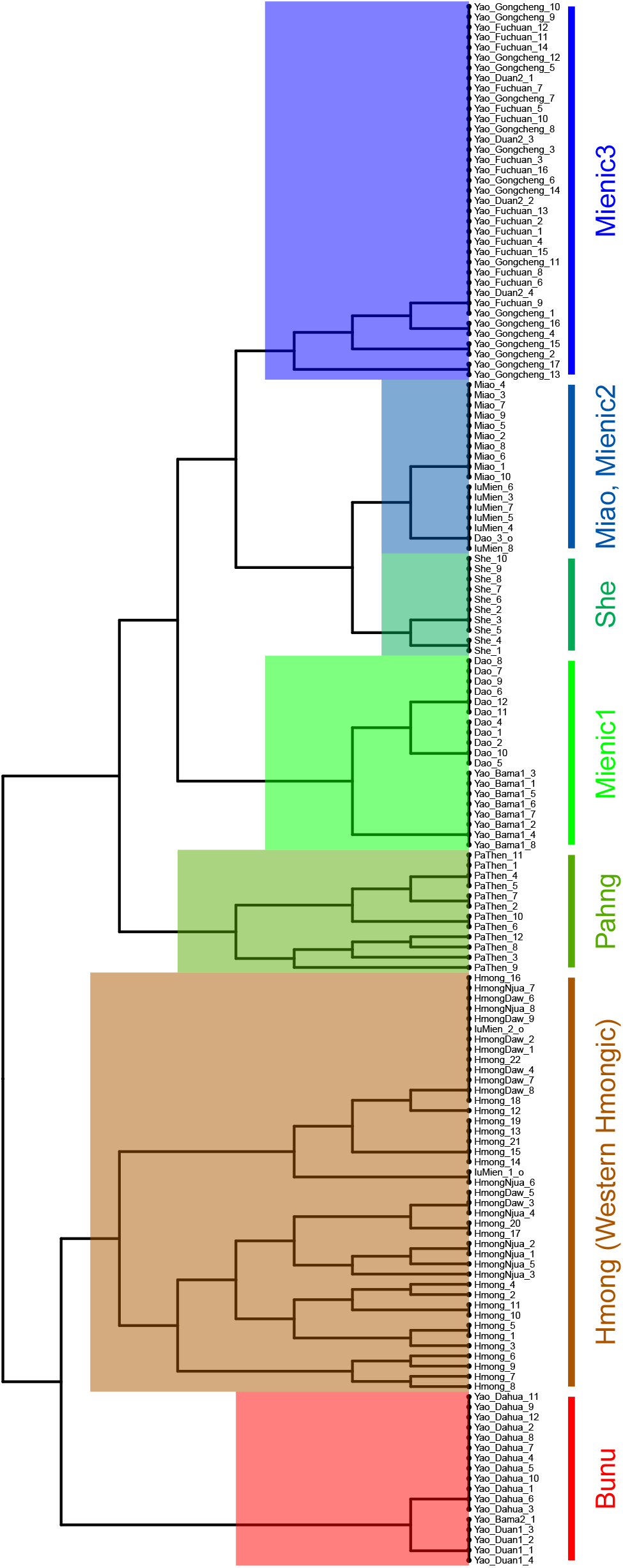
FineSTRUCTURE dendrogram of HM-speaking populations.

The hierarchical structure of genetic clusters partially reflects their linguistic classifications (Figure 3), which supports the parallel between language differentiation and population differentiation in the HM history to some extent. The highest level of the hierarchical tree separates Bunu and Hmong on the left from all the other HM speakers on the right (Figure 3). This indicates that Bunu and Hmong, both of whose languages fall under the classification of ‘West Hmongic’ or ‘Chuanqiandian’ with broad consensus (Strecker, 1987, Ratliff, 2010, Ratliff, 2021, Wang, 1983a) [*Ratliff 2010, p3*], share an extra founder event in relation to other HM speakers, in addition to the founder event shared by all the HM speakers. Given the fact that Bunu and Hmong belong to different officially recognized ethnicities (Yao and Miao, respectively), linguistic affinity is more consistent with the shared genetic history than the classification of ethnicities for both groups. In the right meta-cluster, PaThen from Vietnam (Pahng) splits first from the others (Figure 3), which is in accordance with the fact that they are linguistically most distantly related to other Hmongic languages (Ratliff, 2010, Ratliff, 2021) [*Ratliff 2010, p3*]. Congruent with the fact that the Mienic Kim Mun language is spoken by both Yao_Bama (Figure 1B) and Dao from Vietnam (Scott, 2009) [*Scott 2010, p410*], the majority of both groups are sister clusters in fineSTRUCTURE (‘Mienic1’, Figure 3). By contrast, geography explains less for the genetic structure, as most sister clusters do not share close proximity (e.g., Hmong and Bunu, Dao and Yao_Bama1).

Nevertheless, instead of a sharp split between Hmongic- and Mienic-speaking populations, the genetic clusters with both linguistic affiliations intervene with each other. For example, Mienic-speaking Iu Mien from Thailand forms a sister cluster with North Hmongic-speaking (Ratliff, 2021) Miao (Qo Xong). Therefore, besides genetic isolations, admixture from external groups may also play an extra role in the genetic differentiation among HM-speaking populations. To increase the power of ancestral history inference, we separated all the HM-speaking individuals with the same ethnic labels but different fineSTRUCTURE clusters into distinct populations for all the following analyses, following López *et al*. (2021) (see Method).

### Genetic origin of the ancient GaoHuaHua population

Since the ancient HM-related GaoHuaHua genomes are pseudo-haploid, we performed allele-frequency-based analyses to determine their genetic affinity to all the present-day HM-speaking populations (Figure 4).

**Figure 4.**
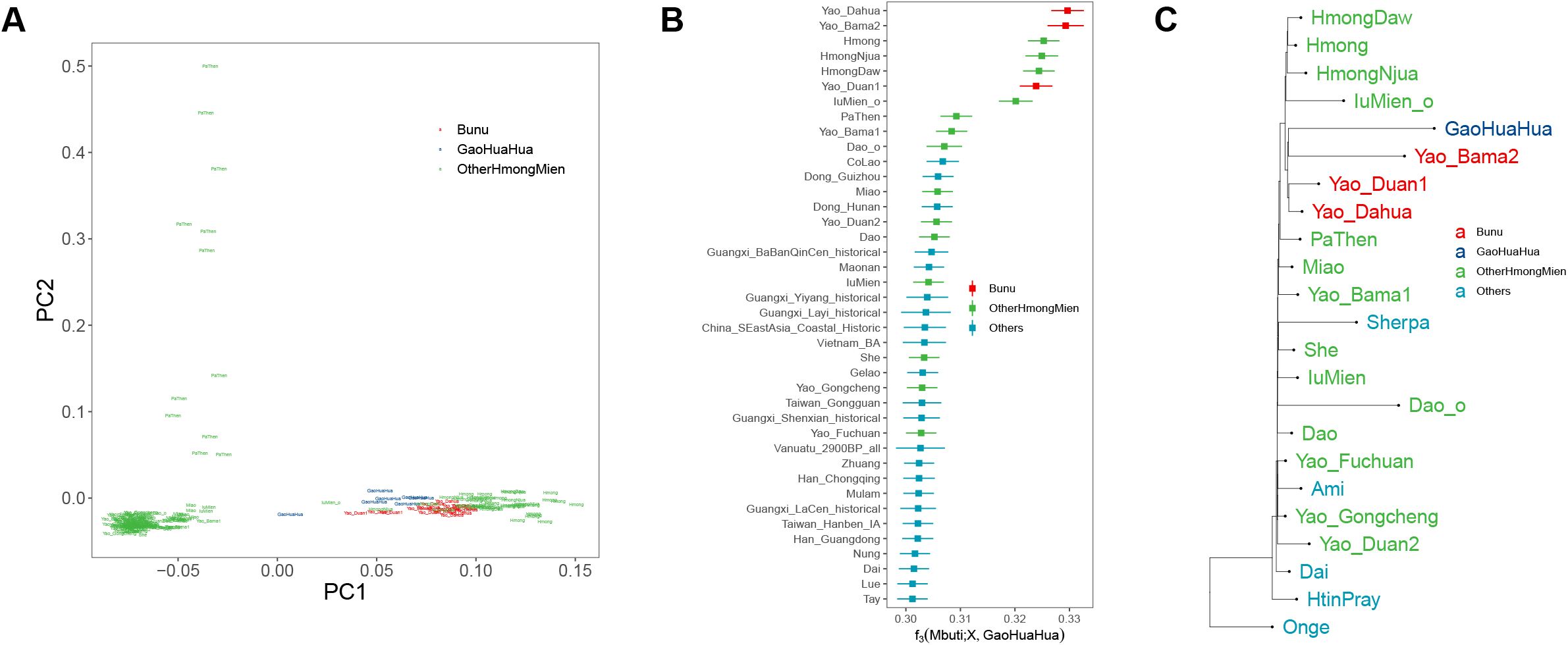
Genetic relationship of GaoHuaHua ancient samples with present-day HM speakers. (A) PCA exclusively for GaoHuaHua and modern HM speakers. (B) Outgroup *f_3_*(Mbuti; X, GaoHuaHua) where X are all the East Asian populations. (C) TreeMix phylogeny with GaoHuaHua and modern HM speakers.

In the PCA exclusively for GaoHuaHua and present-day HM-speaking populations (Figure 4A), PC1 separates West Hmongic-speaking Bunu and Hmong from all the other HM-speaking populations, while PC2 further makes PaThen stands out. GaoHuaHua individuals are projected together with West Hmongic speakers, particularly Bunu-speaking Yao_Duan1, Yao_Dahua, and Yao_Bama2. A consistent pattern can be observed in outgroup-*f_3_* (Figure 4B), where two of the three Bunu-speaking populations share the most genetic drift with GaoHuaHua (Yao_Dahua and Yao_Bama2). Then, we applied TreeMix to generate a maximum likelihood tree with no admixture event and rooted in Onge (Figure 4C). According to the phylogeny, Hmong speakers (Hmong, Hmong Daw, Hmong Njua) and Bunu speakers (Yao_Dahua, Yao_Duan1, Yao_Bama3) are sister clades, whereas GaoHuaHua belongs to the clade of Bunu speakers. Therefore, we conclude that GaoHuaHua is genetically closer related to Bunu speakers than to any other present-day HM-speaking populations. The inferred genetic connection between GaoHuaHua and Bunu speakers is congruent with the fact that the funeral practices in GaoHuaHua sites are similar to present-day Baiku Yao (Wang *et al*., 2021b, Zhang *et al*., 1986), a Bunu-speaking group (Meng, 2001) [*pĵ*]. GaoHuaHua also provides a time calibration for serial founder events in the population history of HM speakers, as the divergence of the Hmong clade and the Bunu clade (Figure 4C) must precede the date of GaoHuaHua (1437–1656 CE) (Wang *et al*., 2021b).

### Demographic dynamics of HM population history

The distribution of shared haploblock in different lengths can provide pivotal insights into the pattern of effective population size (*N_e_*) in population history, i.e., demographic history (Ceballos *et al*., 2018, Ralph and Coop, 2013, Browning and Browning, 2013a, Palamara *et al*., 2012, Ringbauer *et al*., 2021). We sought to characterize the demographic history of HM speakers by using the pattern of shared haploblocks in three different levels: (1) parental relatedness within each individual; (2) individuals within a population; (3) in pairwise populations.

Runs of homozygosity (ROH) is a commonly used measurement for how the parents of an individual are genetically related to each other (Ceballos *et al*., 2018, Ringbauer *et al*., 2021). Theoretically, consanguineous marriage between relatives who share a common ancestor within a few generations tends to result in an excessive number of long ROHs. By contrast, a rapid decline of *N_e_* (i.e., genetic bottleneck) tends to result in the sharing of relatively few ancestors in long-term history and an excessive number of short ROHs (Ceballos *et al*., 2018). As instructed by Ceballos *et al*. (2018), we computed the average ROH of HM-speaking and other South Chinese populations (Figure 5A).

**Figure 5.**
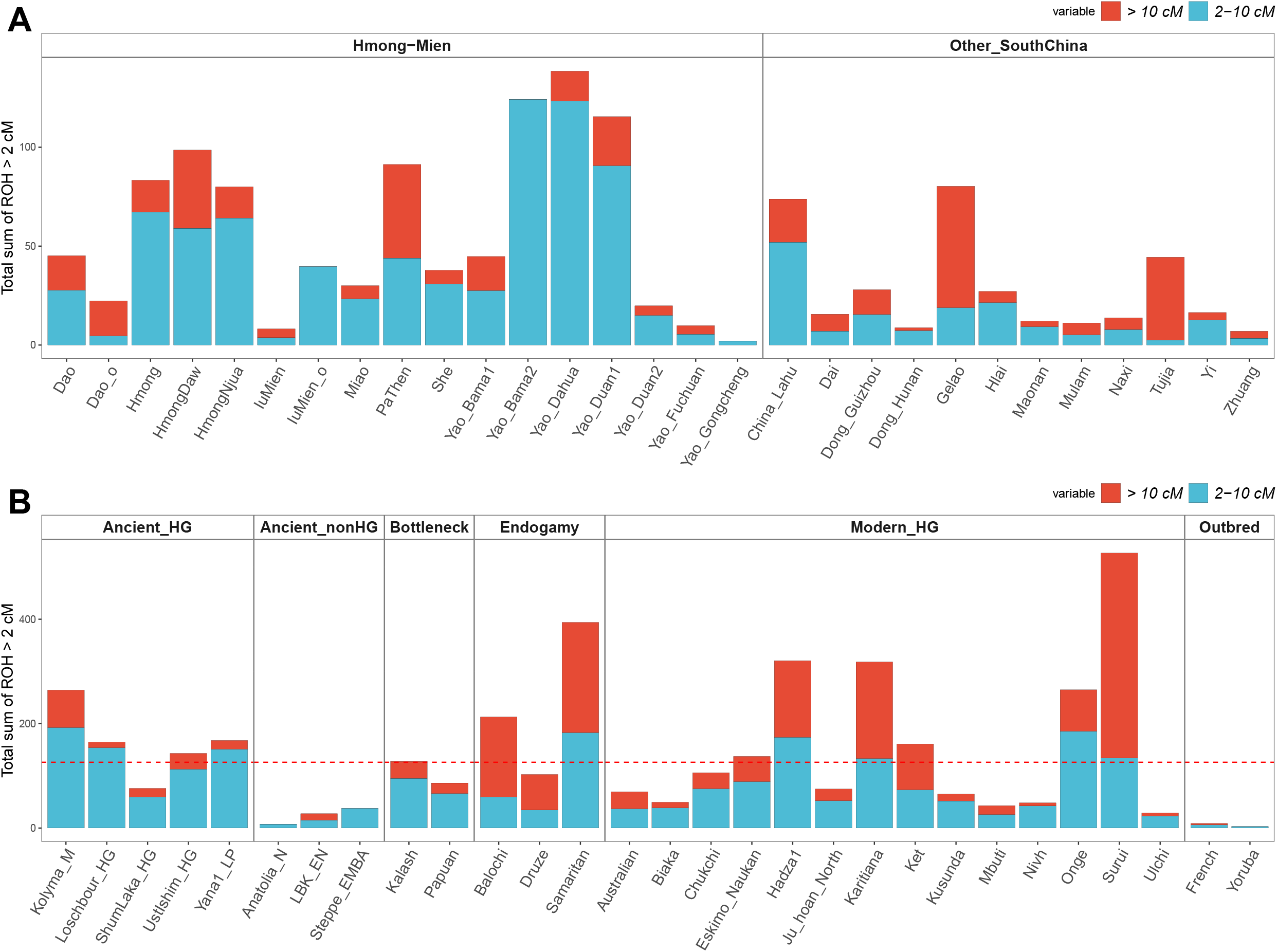
ROH analysis. (A) ROH of HM speakers and other South Chinese populations. (B) ROH of selected worldwide ancient and present-day populations. Dash line shows the average total sum of ROH > 2 cM for all the three Bunu populations (Yao_Dahua, Yao_Duan1, and Yao_Bama2).

Compared with non-HM-speaking populations in South China, especially Kra-Dai-speaking sedentary lowland wet-rice farmers (e.g., Zhuang and Dai), most of the HM-speaking populations have an excessive level of the total length of ROH (Figure 5A). The majority of HM-speaking populations, except for Dao_o and Pa Then, have an enrichment of short ROHs (2–10 cM) than long ROHs (> 10 cM), which indicates that their excessive ROH level is primarily a result of strong genetic bottlenecks in their population history rather than recent consanguineous marriage. Notably, the three Bunu-speaking populations (Yao_Dahua, Yao_Duan1, and Yao_Bama2) have the strongest level of ROH among all the Southern Chinese populations, which is even stronger than their sister clade, Hmong-speaking populations (Hmong, Hmong Daw, Hmong Njua).

We compared the average ROH of Bunu-speaking populations (126.1 cM, dash line, Figure 5B) in relation to chosen ancient and modern global populations with diverse modes of subsistence and marital practice (Figure 5B). In modern populations, the ROH level of Bunu speakers is comparable to Kalash, who is known for their high degree of genetic isolation (Ayub *et al*., 2015, Hellenthal *et al*., 2016), and even stronger than many hunter-gatherer populations in Africa (Mbuti, Biaka, and Jul’hoan) and Asia (Chukchi, Eskimo, Kusunda, Nivh, and Ulchi). As for the ancient genomes, the total ROH of ancient hunter-gatherers is ~ 0.6 – 2.1 folds as Bunu speakers, while one of Bunu speakers is 2–16-fold higher than ancient plant farmers (Anatolia_N, LBK_EN) and pastoralists (Steppe_EMBA). In conclusion, the extent of genetic isolation and bottleneck in Bunu speakers is more similar to the one of ancient and modern hunter-gatherers rather than neighboring sedentary lowland wet-rice farmers, which is likely due to their practice of slash-and-burn agriculture during their history.

We applied *IBD-N_e_* to infer the change of Ne in HM-speaking populations over time from the distribution of identity-by-descent (IBD) (Figure 6A) (Browning and Browning, 2015), and we observed bottlenecks taking place in most of the HM-speaking populations. Except for Hmong Njua (1303 CE), the bottleneck time estimates for all the Bunu-speaking populations [Yao_Duan1, 1037 CE; Yao_Dahua, 1079 CE] and Hmong-speaking populations [Hmong, 1107 CE; Hmong Daw, 1107 CE] are very close to each other. Since such a time range of bottlenecks obviously predates the time of Bunu-related GaoHuaHua (1437–1656 CE) (Wang *et al*., 2021b), this might reflect a shared bottleneck by all the West Hmongic speakers. Bottleneck events occurring around the South Song Dynasty (1127–1279 CE) are inferred in Miao (1275 CE) and Yao_Bama1 (1247 CE). The bottleneck time estimate for She (1471 CE) postdates the first occurrence of She in Fujian (1236 CE) (Chan, 2006), suggesting that the bottleneck of She might occur after their initial settlement in Fujian. More recent estimates of bottleneck events are observed in HM-speaking populations in Vietnam, i.e., Pa Then (1639 CE), Dao (1667 CE), as well as a second bottleneck for Hmong (1779 CE), which indicates the occurrence of founder events along with their migration from South China to Vietnam.

**Figure 6.**
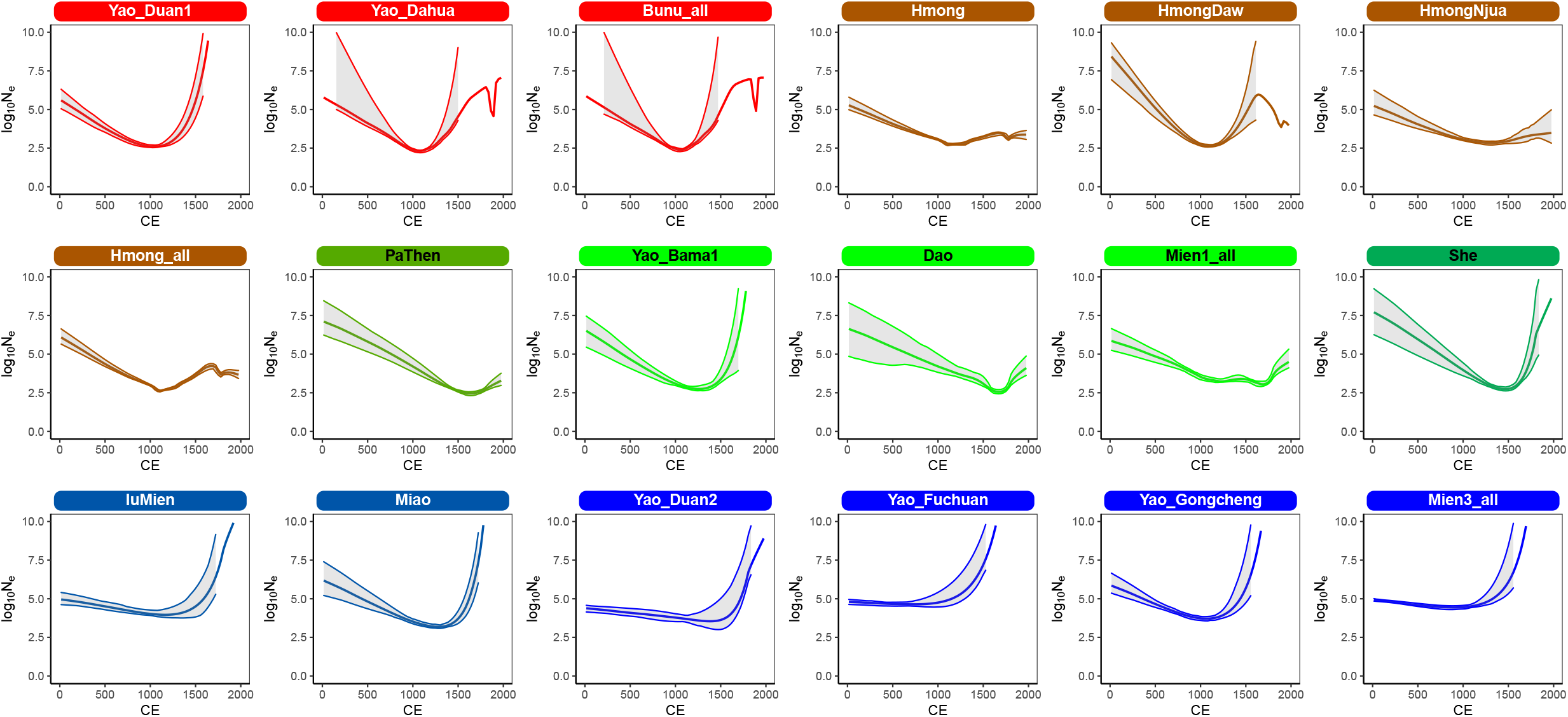
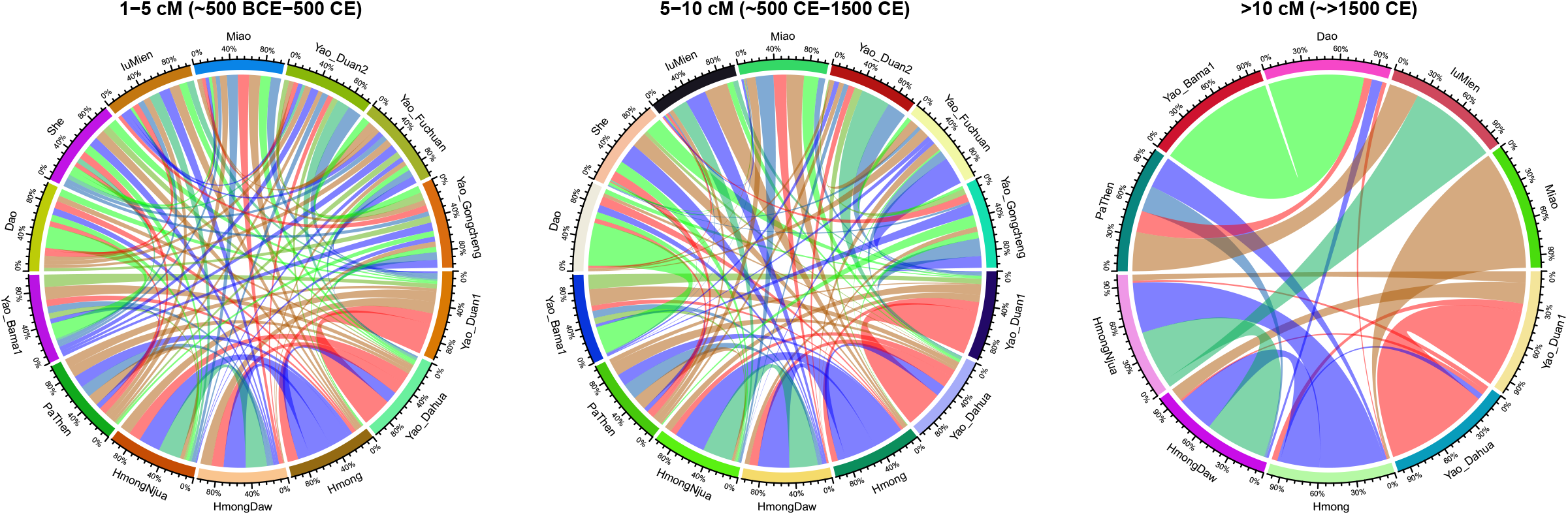
IBD analysis. (A) The trajectories of *N_e_* of HM-speaking populations estimated from intra-population IBD. (B) Inter-population IBD sharing of HM speakers.

We then focused on the intra-population sharing of IBD segments in HM speakers (Figure 6B). Following the instruction by Ralph and Coop (2013), we classified the IBD segments shared by pairwise populations into three categories: 1–5 cM, 5–10 cM, and > 10 cM, which approximately correspond to the time range of 500 BCE – 500 CE, 500 CE – 1500 CE, and > 1500 CE, respectively. In ~500 BCE – 500 CE, each HM-speaking population tends to share common ancestors with all the other HM speakers. In ~500 CE – 1500 CE, HM-speaking populations tend to share more common ancestors with closely related clusters identified in fineSTRUCTURE, which roughly overlaps the frequent emergence of various ethnic groups related to present-day HM speakers (e.g., She and Yao) (Zeng, 2005, Scott, 2009, Litzinger, 1995). Since ~1500 CE, the majority of shared common ancestors are within the sister clusters in fineSTRUCTURE.

### Admixture scenario of HM population history

Besides the variation of *N_e_* over time, admixture with external populations is another factor that can contribute to the genetic differentiation among the HM-speaking populations. To this end, we verified and inferred the temporal scenario of admixture events in HM population history using the haplotype-based fastGLOBETROTTER (Hellenthal *et al*., 2014, Wangkumhang *et al*., 2022). We used all three populations in the ‘Hmong’ cluster defined by fineSTRUCTURE (i.e., Hmong, Hmong Daw, and Hmong Njua, Figure 3, referred to as ‘Hmong_all’) to surrogate the shared ancestry by HM speakers, along with 120 ancient and modern global populations to surrogate other ancestries potentially contributing to present-day HM speakers genetically (SI Table 2). In addition to the other 14 HM-speaking populations, we also included 10 non-HM populations who likely received recent HM-related gene flow (SI Table 2) with the HM-related component > 5% in ADMIXTURE when K = 10 (SI Figure 1).

In all the 24 target populations, 16 of them show sufficiently strong statistical evidence for ‘one-date’ admixture between two primary ancestral sources (fit quality for one event ≥ 0.990, fit event quality for two events ≥ 0.999, goodness-of-fit *r^2^* = 0.316–0.859 for single admixture, Figure 6 and SI Table 3). In contrast, HM-speaking Dao_o, Yao_Duan2, Yao_Fuchuan, and Yao_Gongcheng do not show significant statistical evidence for recent admixture (‘best guess conclusion’ inferred as ‘unclear signal’, SI Table 3). Regarding the admixing sources, one of the primary sources is always inferred to be HM-like (Hmong Daw), while the other source is either Kra-Dai- (Hlai) or Southern Han Chinese-related (Han_Guangdong, Han_Fujian, or Han_Hubei, Figure 7B & SI Table 3). This can be explained by the fact that all the non-HM-speaking speakers inferred with this HM-related admixture are either Kra-Dai speakers affiliating to Kra (Gelao/Cò’ Lao) or Kam-Sui (Dong_Hunan, Dong_Guizhou, and Maonan) or Tujia, who is historically Tibeto-Burman-speaking but largely Sinicized. The inferred average admixture dates for most of the populations (Figure 7A) overlap the more frequent presence of Miao and Yao in Chinese historical documents since the Ming Dynasty (1368–1644 CE) (Diamond, 1995, Faure, 2006). The inferred dates of admixture events in Bunu-speaking Yao_Dahua (997 CE, 95% confidence interval (CI) 791–1223 CE), Yao_Duan1 (1108 CE, 95% CI 832–1345 CE), and Yao_Bama2 (1334 CE, 95% CI 1089 – 1594 CE, Figure 7A) suggest that the genetic profile of Bunu speakers was largely fixed prior to or at the time of GaoHuaHua (1437–1656 CE). The inferred dates of IuMien (1826 CE, 95% CI 1538–1947 CE) and IuMien_o (1745 CE, 95% CI 1644 – 1866 CE) roughly coincide with their migration from South China to Southeast Asia.

**Figure 7.**
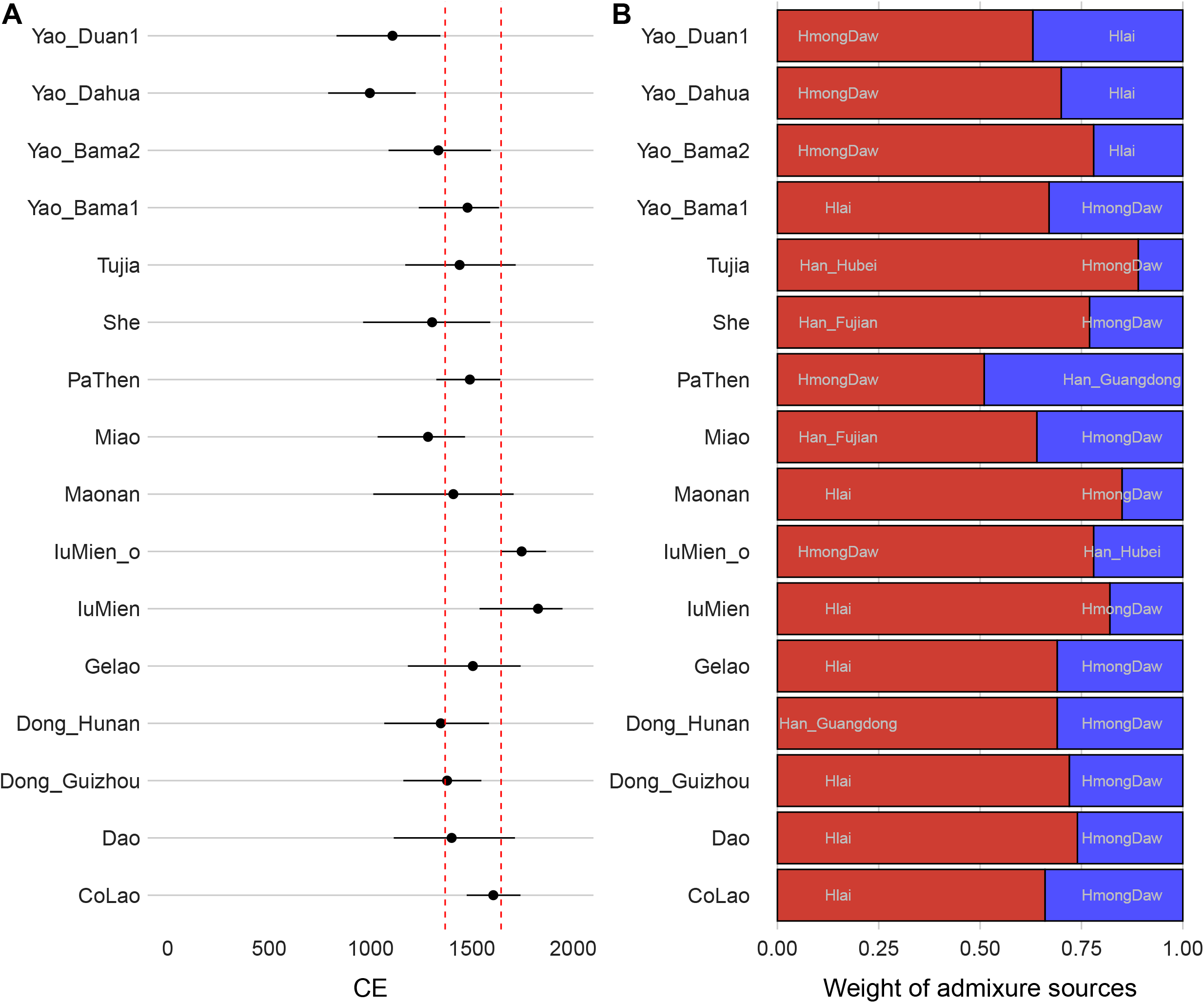
Admixture dates and most likely contributors inferred by fastGLOBETROTTER. (A) Point estimates and 95% CIs of inferred admixture dates in HM speakers and neighboring populations. (B) Contributions of two-way admixture and single best-matching proxies for each ancestry source.

To further characterize a global picture of the fine-scale ancestry composition in HM speakers, we applied SOURCEFIND (Chacón-Duque *et al*., 2018) with the same target and surrogate sets as fastGLOBETROTTER analysis (SI Table 2). The variation in the proportion of the HM-like ancestry (‘Hmong_all’, Figure 8A) in HM-speaking populations majorly corresponds to their genetic clustering in fineSTRUCTURE (Figure 3): IuMien_o (81.5%), Bunu-speaking Yao_Dahua (67.9%), Yao_Bama2 (66.7%), and Yao_Duan1 (58.3%) harbour the largest proportion of this HM-like source, which is followed by PaThen (26.0%). Notably, Kra-speaking Cò’ Lao (16.6%) and Gelao (9.7%), as well as Kam-Sui-speaking Dong_Guizhou (9.6%) and Dong_Hunan (11.6%), possess a comparable level of this HM-like ancestry as many HM speakers. Rather than a term specific HM speakers, the exonym ‘Miao’ is more likely to be a collective term for indigenous groups in Southwest China (or more exactly, in ‘Miao territory’/Miao Jiang (苗疆) of Guizhou and western Hunan) who were out of the direct administration during Ming and Qing dynasties, which also includes Kra-Dai-speaking populations in this region. (Diamond, 1995, Scott, 2009) [*p110, p121*]. The inferred admixture dates of these populations (1346 – 1606 CE, Figure 7A) roughly fall within the reign of the Ming dynasty (1368 – 1644 CE), which suggests that massive admixture forming the ‘ HM Cline’ described previously (Huang *et al*., 2022) were among ethnic groups who were linguistically heterogeneous but with similar ethnic identity during this period.

**Figure 8.**
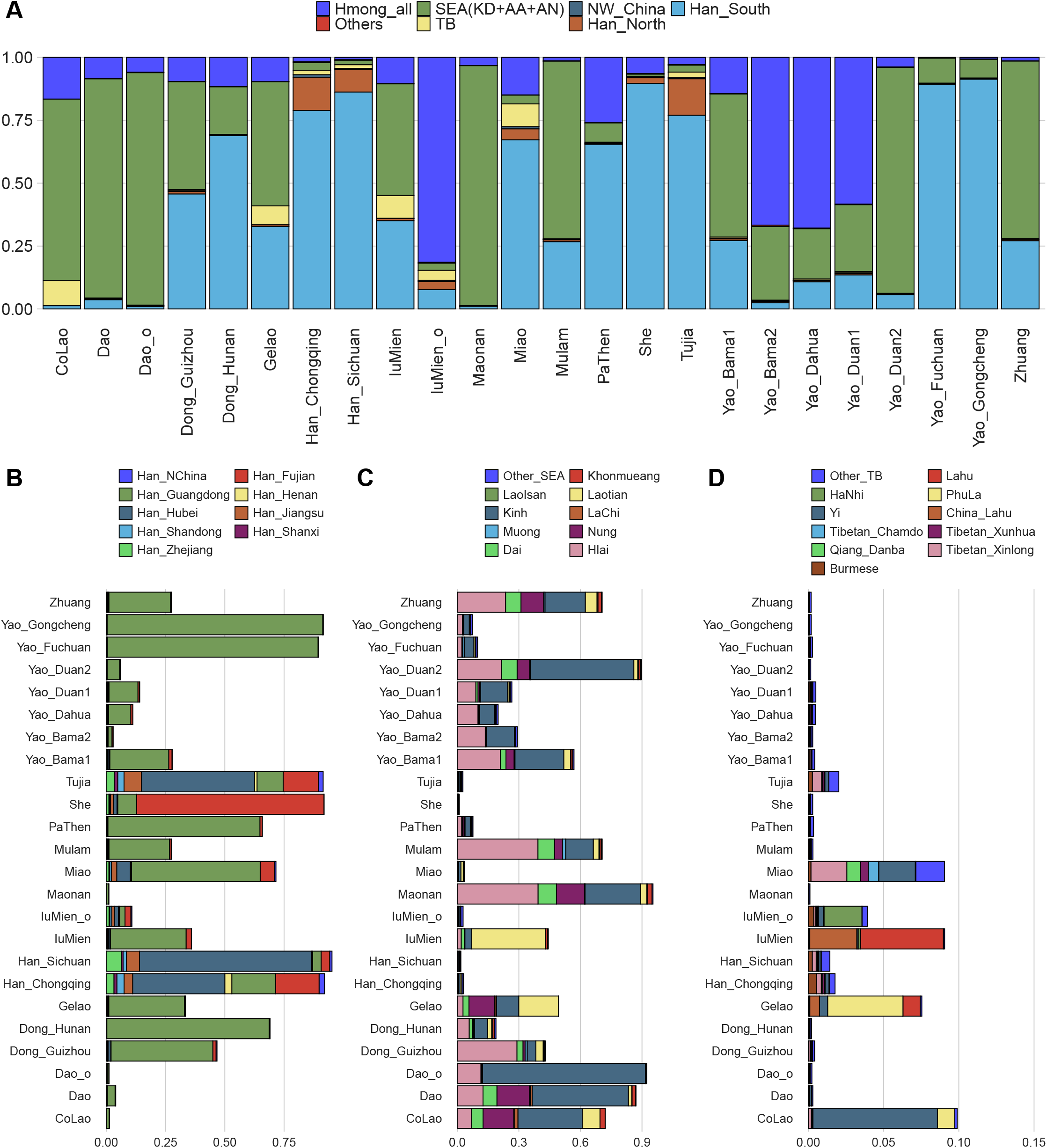
Finescale ancestry composition inferred by SOURCEFIND. (A) A summary of language family-level ancestry contribution. Ancestry contribution in detail from proxies of (B) Han Chinese, (C) Southeast Asians (Kra-Dai, Austronesian, and Austroasiatic speakers), and (D) Tibeto-Burman speakers.

Consistent with the pattern inferred by fastGLOBETROTTER, the majority of non-HM-like ancestry in HM speakers comes from either southern Han Chinese-like sources (Figure 8A), primarily surrogated by Han_Guangdong (Figure 8B), or Southeast Asian-like sources (Figure 8A), primarily surrogated by Kinh, Hlai, and Nung (Figure 8C). However, the distribution of these non-HM-like ancestries in HM-speaking populations, especially those in South China, varies geographically (Figure 8A). All the Yao populations from localities in western Guangxi (i.e., Duan, Dahua, and Bama, Figure 1A) derive more of their non-HM-like ancestry from Southeast Asian-like sources than southern Han Chinese-like ones (Figure 8A). By contrast, for HM-speaking populations in eastern Guangxi (Yao_Gongcheng and Yao_Fuchuan, Figure 1A), eastern Guizhou (Miao, 28N, 109E), and northeast Fujian (She, 27N, 119E) (Cann *et al*., 2002), their non-HM-like ancestry is primarily surrogated by southern Han Chinese-like sources (Figure 7A & B). Miao, Iu Mien, Gelao, and Cò’ Lao derive additional Tibeto-Burman-like ancestry (> 5%, Figure 8A) primarily surrogated by Loloish-speaking Yi, Phù Lá, and Lahu.

We also observed special admixture histories in some HM-speaking populations. Congruent with their admixture date interval (1304 CE, 95% CI 963 – 1591 CE, Figure 7A), the largest proportion of ancestry surrogated by Han_Fujian (79.0%, Figure 8B) in She confirms that they acquired the majority of their southern Han Chinese-related ancestry only after their settlement in Fujian. Likewise, the extra genetic influx surrogated by Laos (36.0%, Figure 8C) is observed in Iu Mien currently living in Northeast Thailand (Kutanan *et al*., 2021), which is compatible with the scenario of admixture during their migration out of South China. Conforming the results from fastGLOBETROTTER (SI Table 3), Yao_Fuchuan and Yao_Gongcheng – both of whom include both Iu Mien speakers and Pingdi Yao (Figure 1B) – have a negligible amount of HM-like ancestry (0.2% and 0.6%, Figure 8A) and a small amount of Southeast Asian-like ancestry (9.9% and 7.4%, Figure 8A), but they derive the majority of their ancestry surrogated by Han_Guangdong (88.9% and 90.9%, Figure 8B). Since SOURCEFIND infers a considerable amount of Southeast Asian-like ancestry in all the Kra-Dai-speaking populations (18.9 – 95.4%, Figure 8A), it is unlikely that all of the Han_Guangdong-related ancestry in both groups derive from Kra-Dai speakers, while at least some come from southern Han Chinese. Our result is compatible with the theory that some of the Yao people may have originated from Han Chinese who avoided tax and corvée and adopted the ethnic identity of ‘Yao’ (Scott, 2009) [*p121, p125*].

## Discussion

By leveraging newly reported genome-wide data and haplotype-based methods, our study extends the understanding of how historical isolation and admixture events shape the genetic structure of HM speakers to an unprecedented level of resolution. For example, we are able to identify the closely related genetic clusters in HM speakers (e.g., Yao_Bama1 and Dao, Figure 3) and the external gene flow from Han Chinese in a province level (e.g., Han_Guangdong vs Han_Fujian, Figure 8). The genetic structure in such a fine scale enables us to discuss further the genetic impact of the unique mode of subsistence that makes HM speakers distinctive from most of the other southern East Asian groups in history: slash-and-burn and shifting agriculture.

The practice of slash-and-burn agriculture tends to be associated with reduced population sizes. In South China and Mainland Southeast Asia, wet-rice agriculture is majorly practiced in lowland valleys where most of the easily cultivable land is distributed, so it promises high return per unit of land and high population density (Scott, 2009) [*p13, p41, p74*]. By contrast, slash-and-burn agriculture is more commonly practiced in mountainous areas because there is much sparser arable land, and the slash-and-burn technique transforms forests into nutrient-rich soil suitable for farming (Scott, 2009) [*p18*]. Consequently, the population density for slash-and-burn farmers is much lower than for wet-rice farmers, approximately an order of magnitude (Bellwood, 1993). Therefore, our observation of the strong genetic bottleneck in HM speakers – especially Bunu, Hmong, and Pahng/Pa Then (Figures 5 & 6) – supports the theoretically expected demographic patterns of slash-and-burn farmers.

Since slash-and-burn soils lose fertility quickly, slash-and-burn farmers must regularly move to a new place and reclaim a new land using the slash-and-burn technique, i.e., shifting farming. Consequently, slash-and-burn farmers, e.g., HM speakers, tend to be more nomadic than sedentary wet-rice farmers. This explains why the genetic substructure of HM speakers is less geographically associated (Figures 2 & 3) in contrast to an overall geographically associated genetic structure in East Asia (Wang *et al*., 2021a). By contrast, the distribution of external ancestries in HM speakers is more geographically associated (Figure 8), suggesting the pattern of admixture after settlement. The long-term genetic bottleneck and nomadic lifestyle also led to serial founder events and population differentiation that parallel language differentiation in West Hmongic speakers (Figure 3). Our analysis highlights that the practice of slash-and-burn and shifting agriculture is a crucial factor associated with the genetic and demographic history of HM speakers, which has rarely been addressed in previous genetic studies about HM-speaking populations.

On a worldwide scale, previous genetic studies have revealed the impact of geographic isolation, modes of subsistence, and marital practice on effective population size (Ceballos *et al*., 2018, Ringbauer *et al*., 2021, Tournebize *et al*., 2022). Regarding modes of subsistence, plant farmers tend to have a lower degree of a founder effect, while a nomadic lifestyle leads to enhanced genetic bottleneck (Tournebize *et al*., 2022). However, it remains unclear whether plant agriculture or nomadism is the more decisive factor in terms of the effective population size in previous studies. Therefore, the evidence from HM speakers – who are farmers but historically nomadic – supports that nomadism plays a more crucial role in demographic patterns.

In conclusion, our study reveals the recent isolation and admixture events within the recent ~2,500 years that contribute to the fine-scale genetic formation of present-day HM-speaking populations and investigates the impact of sociological/anthropological factors – especially slash-and-burn and shifting agriculture – on the genetic pattern of HM speakers. Given the power of GaoHuaHua in the calibration of HM history, we predict that more ancient genomes from Southwest China in the recent ~2,500 years will shed new light on the origin and differentiation of HM-speaking populations. Our study also highlights the importance of incorporating currently underrepresented groups in future genetic studies, e.g., genome-wide association studies (GWAS).

## Methods

### Sample collection and genomic data curation

We collected saliva and blood samples from 67 Yao individuals from five autonomous counties (Du’an, Dahua, Bama, Gongcheng, Fuchuan) in Guangxi, China (SI Table 1). All sample donors read and signed informed content, and this research was approved by the Ethical Committee of Xiamen University (approval number: XDYX2019009). All of the processes were in accordance with the corresponding ethical principles. Then, we obtained the genotyped data of these samples using an Infinium Global Screening Array covering 699,537 genome-wide SNPs. We imputed the genotype using Eagle v2.4.1 (Loh *et al*., 2016) and Minimac4 (Das *et al*., 2016) with default parameters (with a chunk size of 10 Mb and step size of 3 Mb) against the 1000 Genomes project Phase3 v5 reference haplotypes (Consortium *et al*., 2015). We then removed SNPs with imputation quality < 0.3, MAF < 1% or missing rate > 2%. To identify presumably related individuals, we used PLINK v1.9 (Purcell *et al*., 2007) with the parameter ‘-genome’ to estimate PI_HAT values for each pair of newly sampled individuals and removed all three individuals with PI_HAT > 0.15 in all the subsequent analyses. After that, we merged our data with ‘1240k+HO’ dataset in Allen Ancient DNA Resource (AADR) V50.0 (Reich, 2022) and recently published modern (Liu *et al*., 2020, Kutanan *et al*., 2021) and ancient genomes (Wang *et al*., 2021b, Liu *et al*., 2022) for co-analysis, resulting in 362,468 autosomal SNPs used in all the following analyses. We only kept diploid modern and ancient genomes for haplotype-based analyses and added ancient diploid genomes (Devil’s Gate, Yana, and Kolyma) from Sikora *et al*. (2019). We identified outliers given the results from PCA (Patterson *et al*., 2006) and fineSTRUCTURE (Lawson *et al*., 2012). Non-HM outliers were removed in subsequent analysis, while HM outliers were marked with labels (‘_o’ or ‘_2’).

### Allele frequency-based analyses

The program *smartpca* packed in EIGENSOFT (Patterson *et al*., 2006) was used to perform PCA with default parameters and lsqproject: YES, numoutlieriter: 0. Only modern samples were used to construct principal components, whereas ancient samples were projected. For ADMIXTURE analysis (Alexander *et al*., 2009), we first used PLINK v1.9 (Purcell *et al*., 2007) to prune linkage disequilibrium (-indep-pairwise 200 20 0.4). ADMIXTURE v1.3.0 was then performed from K=2 – 20 (SI Figure 1) with unsupervised mode and default parameters. K = 10 was reported for its lowest cross-validation error. We used ADMIXTOOLS (Patterson *et al*., 2012) to compute outgroup *f_3_*-statistics with Mbuti as the outgroup for East Asians. We used *TreeMix* (Pickrell and Pritchard, 2012) to generate allele frequency-based phylogeny with no migration (-m 0), a window size of every 500 SNPs (-k 500), and Onge as the root.

### Haplotype-based analyses

We used SHAPEIT v2 (O’Connell *et al*., 2014) to phase diploid ancient and modern genomes in our data. According to López *et al*. (2021), we used the following procedures to generate a copying-vector profile using ChromoPainter (Lawson *et al*., 2012). We first estimated the parameters for mutation/emission (*Mut*, ‘-M’) and switch rate (*N_e_*, ‘-n’) with ten steps of the Expectation-Maximization (E-M) algorithm for chromosomes 1, 8, 15, and 22 for the first 10 individuals. Then, we used the fixed *Mut* and *N_e_* values estimated in the previous procedure to run ChromoPainter for all the individuals. For the copying-vector profile used for fineSTRUCTURE (Lawson *et al*., 2018) analysis, we used only HM-speaking individuals as both donors and recipients with the fineSTRUCTURE normalization parameter ‘c’ estimated to be 0.395. Then, we used fineSTRUCTURE to perform 200,000 sample iterations under Markov chain Monte Carlo (MCMC) with the first 100,000 samples as burn-ins, and the inferred trees were sampled every 1,000 iterations. After that, we carried out 100,000 hill-climbing steps, and fineSTRUCTURE summarized the most likely tree.

For SOURCEFIND (Chacón-Duque *et al*., 2018) and fastGLOBETROTTER (Hellenthal *et al*., 2014, Wangkumhang *et al*., 2022) analyses, we used all the three populations in the ‘Hmong’ cluster defined by fineSTRUCTURE (i.e., Hmong, Hmong Daw, and Hmong Njua, Figure 3, referred as ‘Hmong_all’) to surrogate the shared ancestry by HM speakers, along with 120 ancient and modern global populations to surrogate other ancestries potentially contributing to present-day HM speakers genetically (SI Table 2). In addition to the other 14 HM-speaking populations, we also included 10 non-HM populations who likely received recent HM-related gene flow (SI Table 2) with the HM-related component > 5% in ADMIXTURE when K = 10 (SI Figure 1). We generated the copying-vector profiles for both analyses following the suggestions by Wangkumhang *et al*. (2022). All the other parameters of both methods were kept by default. For ROH analysis, we used PLINK v1.9 (Purcell *et al*., 2007) with the parameters following the suggestions by Ceballos *et al*. (2018). We used Refine IBD to obtain pairwise IBD of HM-speaking individuals with default parameters (Browning and Browning, 2013b). For intra-population IBDs, we used all the IBD segments > 1.0 cM to estimate the change of *N_e_* over time using IBDNe (Browning and Browning, 2015) for the recent ~2,000 years. Following Ralph and Coop (2013), we grouped inter-population IBDs into three categories: 1–5 cM, 5–10 cM, and > 10 cM, which approximately correspond to the time range of 500 BCE – 500 CE, 500 CE – 1500 CE, and > 1500 CE, respectively. For fastGLOBETROTTER and IBDNe, we used 1975 CE as the average birthdate for all the modern individuals and 28 years per generation to covert generation into year following López *et al*. (2021).

## ACKNOWLEDGEMENTS

The work was funded by Guangxi First-class Discipline Project for Basic medicine Sciences (No. GXFCDP-BMS-2018), the National Natural Science Foundation of China (32270667), Basic Medical Science and Technology Innovation Education Fund of Guangxi Medical University (No.: GXMUBMSTCF-G10), the “Double First Class University Plan” key construction project of Xiamen University (0310/X2106027), Nanqiang Outstanding Young Talents Program of Xiamen University (X2123302), the Major Project of National Social Science Foundation of China granted to Chuan-Chao Wang (21&ZD285), Major Special Project of Philosophy and Social Sciences Research of the Ministry of Education (2022JZDZ023), and European Research Council (ERC) grant (ERC-2019-ADG-883700-TRAM).

## AUTHOR CONTRIBUTIONS

C.C Wang and Z.Y Xia conducted the project and conceived the idea. Q.Y Deng and X.C Chen collected the samples and carried out the experiments. Z.Y Xia analyzed the data and wrote the paper. C.C Wang edited the draft. All the authors revised the paper.

