## Supplementary figures and images for "Reconstructing the formation of Hmong-Mien genetic fine-structure"

### Supplementary Figure 1

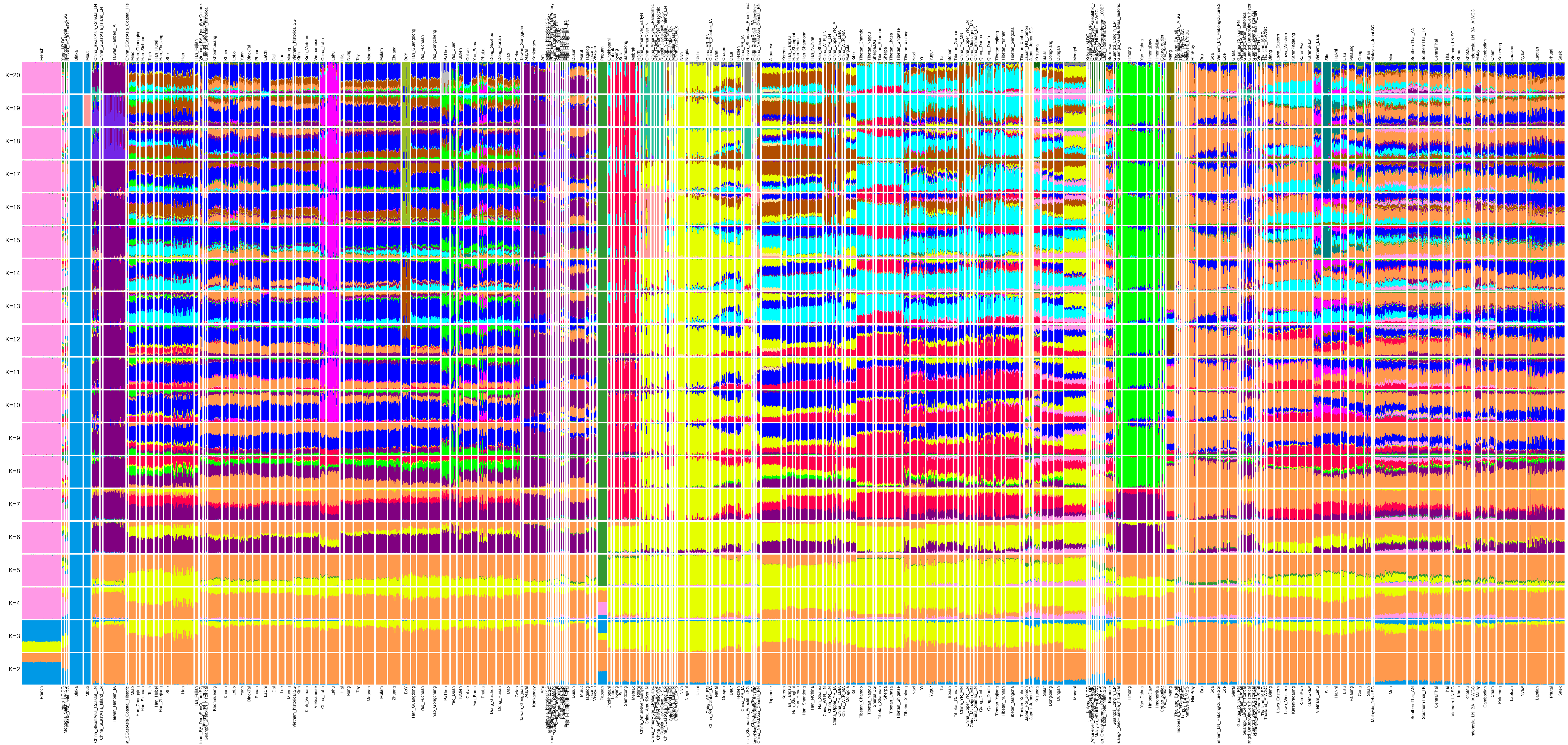
